# The *Chloroflexi* supergroup is metabolically diverse and representatives have novel genes for non-photosynthesis based CO_2_ fixation

**DOI:** 10.1101/2021.08.23.457424

**Authors:** Jacob A. West-Roberts, Paula B. Matheus-Carnevali, Marie Charlotte Schoelmerich, Basem Al-Shayeb, Alex D. Thomas, Allison Sharrar, Christine He, Lin-Xing Chen, Adi Lavy, Ray Keren, Yuki Amano, Jillian F. Banfield

**Affiliations:** University of California – Berkeley; University of California, Berkeley; University of California Berkeley; Sector of Decommissioning and Radioactive Wastes Management, Japan Atomic Energy Agency

## Abstract

The *Chloroflexi* superphylum have been investigated primarily from the perspective of reductive dehalogenation of toxic compounds, anaerobic photosynthesis and wastewater treatment, but remain relatively little studied compared to their close relatives within the larger *Terrabacteria* group, including Cyanobacteria, Actinobacteria, and Firmicutes. Here, we conducted a detailed phylogenetic analysis of the phylum *Chloroflexota,* the phylogenetically proximal candidate phylum *Dormibacteraeota*, and a newly defined sibling phylum proposed in the current study, *Eulabeiota*. These groups routinely root together in phylogenomic analyses, and constitute the *Chloroflexi* supergroup. Chemoautotrophy is widespread in Chloroflexi. Two Form I Rubisco ancestral subtypes that both lack the small subunit are prevalent in *ca. Eulabeiota* and *Chloroflexota*, suggesting that the predominant modern pathway for CO_2_ fixation evolved in these groups. The single subunit Form I Rubiscos are inferred to have evolved prior to oxygenation of the Earth’s atmosphere and now predominantly occur in anaerobes. Prevalent in both *Chloroflexota* and *ca. Eulabeiota* are capacities related to aerobic oxidation of gases, especially CO and H_2_. In fact, aerobic and anaerobic CO dehydrogenases are widespread throughout every class-level lineage, whereas traits such as denitrification and reductive dehalogenation are heterogeneously distributed across the supergroup. Interestingly, some *Chloroflexota* have a novel clade of group 3 NiFe hydrogenases that is phylogenetically distinct from previously reported groups. Overall, the analyses underline the very high level of metabolic diversity in the Chloroflexi supergroup, suggesting the ancestral metabolic platform for this group enabled highly varied adaptation to ecosystems that appeared in the aerobic world.

## Introduction

The phylum *Chloroflexota* is represented by a variety of isolated bacteria and is one of the phyla best studied by classical approaches^[1][2]^. Based primarily on cultivated strains, the phylum Chloroflexi was subdivided into *Anaerolineae, Ardenticatenia, Caldilineae, ca. Thermofonsia, Limnocylindria, Chloroflexia, Dehalococcoidia, Ktedonobacteria, Tepidiformia, Thermoflexia,* and *Thermomicrobia*^[3]^.

The ability to reconstruct draft genomes from metagenomes circumvents the cultivation requirement and has greatly expanded the genomic coverage of the *Chloroflexota* and related bacteria. Two distinct groups represented by genomes from metagenomes are the ANG-CHLX, first reported from soil in northern California^[4]^ and now renamed as Dormibacteraeota^[5]^, and RIF-CHLX, first reported from an aquifer adjacent to the Colorado River near the town of Rifle, CO^[6][34]^ and later designated as *Chloroflexota* class *ca. Limnocylindria*^[7]^. These two groups remain relatively little explored, both from the perspective of their phylogenetic placement and metabolic traits. We use the term C*hloroflexi* supergroup to refer to the *Chloroflexota* and the RIF-CHLX. Whether *Dormibacteraeota* are monophyletic with Chloroflexi and RIF-CHLX and thus part of the supergroup has remained uncertain, although phylogenetic analyses suggest the *Chloroflexi* supergroup and *Dormibacteraeota* are phylogenetically proximal, and so they are included in our analyses.

Within the *Chloroflexi* supergroup are organisms that are obligate H_2_-dependent haloalkane-reducers of the class *Dehalococcoidia*^[8]^, spore-forming members with actinomycetes-like morphology of the class *Ktedonobacteria*^[9][15]^, and the photosynthetic, thermophilic members of the class *Chloroflexia*^[1]^. *Chloroflexus aurantiacus* (class *Chloroflexia*), the type species of the phylum, was first identified from hot springs in 1974^[1]^; their production of bacteriochlorophyll gave them a characteristic green color, thus the name *Chloroflexi*, from greek ‘χλωρός’, meaning ‘green’. Culturing *C. aurantiacus* in the absence of light yields an orange culture, thus the species name *aurantiacus*, from the latin ‘aureum’ for ‘orange’. Newly characterized *Chloroflexota* have been observed which contain Type I photosystem reaction centers^[10]^, although with the exception of these new organisms, all other phototrophic *Chloroflexota* use Type II *pufLM*-type photosystem reaction centers. The majority of identified microorganisms belonging to the phylum *Chloroflexota*, however, lack the capacity to perform photosynthesis. An interesting recent report demonstrated the genomic capacity for production of photosynthetic reaction centers in the *Chloroflexota* order *Aggregatilineales*, within the clade *Anaerolinea*, based on the presence of divergent *pufLM*-like reaction centers^[12]^. Some representatives of the *Chloroflexi* supergroup have the ability for chemoautotrophic fixation of CO_2_^[13]^, oxidation of CO^[14][16]^, and oxidation of H_2_^[16][17]^.

A surprising and interesting recent finding is that bacteria within *Chloroflexota* class *Anaerolinea* have Form I Rubisco sequences which form a clade that is clearly basal to previously known Form I Rubiscos. Form I Rubisco is the enzyme at the heart of the Calvin Benson Bassham (CBB) cycle used by organisms including Cyanobacteria, algae and plants in the fixation of CO_2_. It is considered to be one of the most abundant proteins on the planet, and one of the most important from the perspective of biosphere primary production. Notably, the new clade, referred to as Rubisco Form I’, lacks the small subunit that is required for function of Form I Rubisco. Importantly, the enzyme has been biochemically characterized and its function in the CBB pathway confirmed^[13]^. Based on this research it was suggested that the small subunit evolved to stabilize the octamer when it adapted to an oxygenated atmosphere to increase specificity for CO_2_ over O_2_^[13]^. Form I’ Rubisco has not been found in bacteria outside of the *Anaerolinea*, suggesting that Form I Rubisco evolved within a lineage closely related to the ancestor of the modern *Chloroflexi* from a single subunit form similar to I’. Better understanding of the forms and distributions of Rubisco in the *Chloroflexi* supergroup may provide further insight regarding the evolution of the CBB cycle and clues to the metabolic context into which it evolved.

Here, we assembled a Chloroflexi supergroup genome dataset comprising publicly available and newly reconstructed genomes from groundwater, river sediment and soil environments. We constructed a detailed phylogeny for the supergroup, clarified the relationships among currently described groups and provided a foundation for the analysis of the distribution of metabolic traits across the lineages. Our results highlight several traits differentially distributed among phylum- and class-level lineages within the *Chloroflexi* supergroup and provide the most detailed phylogenetic analysis of the *Chloroflexi* supergroup to date.

## Results

### Ribosomal phylogeny of the Chloroflexi supergroup reveals major subdivisions

We constructed a database that includes 2086 publicly available (see Methods) and 75 newly reconstructed genomes from the *Chloroflexi* supergroup. (**Supplementary Table S1**). A phylogenetic tree based on a concatenation of 16 ribosomal protein sequences from 2069 genomes (**Fig. 1**) reveals multiple potential unstudied class-level lineages within the phylum *Chloroflexota* (**Supplementary data S1**) (**Table 1: Number of genomes per lineage in the***Chloroflexi* **supergroup as assigned by phylogeny**). Proximal to the *Chloroflexota* are two phylum-level lineages, one of which is *ca. Dormibacteraeota.* Some genomes from the second lineage had previously been considered to form a class within *Chloroflexota*, referred to in separate sources as *ellin6529* (GTDB)^[18][19]^, *Edaphomicrobia*^[20]^, RIF-CHLX^[6]^, and *Limnocylindria*^[7]^. For this new and now clearly genomically resolved phylum-level lineage we propose the name *Eulabeiota*, from ancient greek *Eulabeia* (ε⋃λαβεɩɑ, “timidity”, “reverence” or “caution”). Genomes from *ca. Dormibacteraeota* have been recovered from soil and permafrost, whereas genomes from *ca. Eulabeiota* were obtained from diverse environments such as hydrothermal vents, freshwater, groundwater, permafrost, soil, and aquifer sediment.

**Table 1.** Identified clades within the *Chloroflexi* supergroup, including candidate phyla *Dormibacteraeota* and *Eulabeiota*, and the number of identified genomes within each clade.

| Phyletic Group | Number of Genomes |
| --- | --- |
| Anaerolinea | 773 |
| Dehalococcoidia | 551 |
| Candidate phylum Dormibacteraeota | 241 |
| Candidate phylum Eulabeiota | 229 |
| Chloroflexia | 148 |
| Ktedonobacteria | 92 |
| Marine Endosymbiont Lineage | 11 |
| Unresolved lineage (UB5177) | 3 |

**Figure 1.**
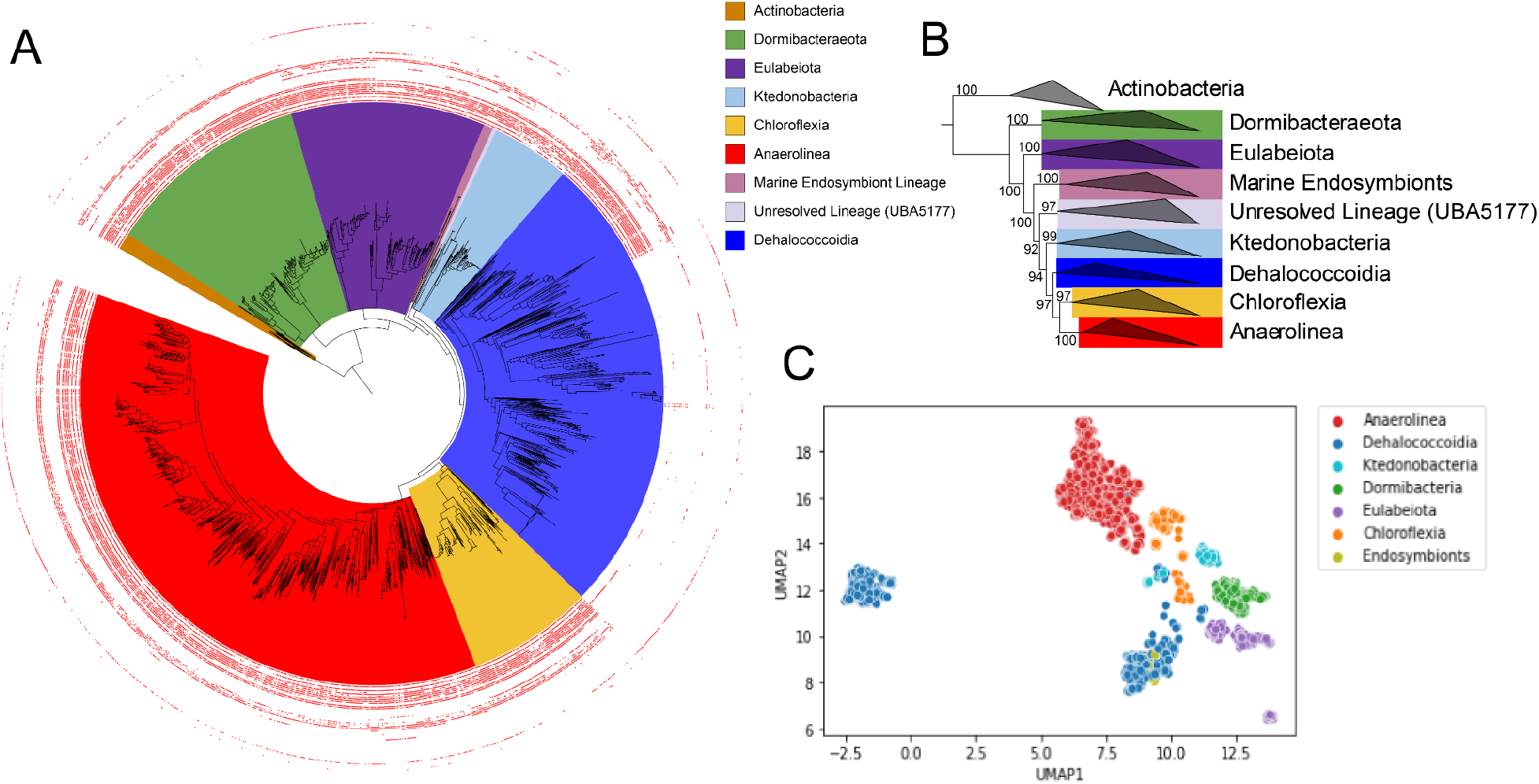
(A) Phylogenetic tree estimated using the PMSF C20 mixture model from concatenated sequences of 16 ribosomal proteins. Shown are the *Chloroflexi* supergroup, including the candidate phyla *Dormibacteraeota* and *Eulabeiota,* and the *Actinobacteria* as an outgroup. Red decorations along the outer ring indicate hits to KOFAM HMMs corresponding to the peptidoglycan biosynthesis pathway (map00550). (B) Rectangular view of the same tree showing the relative positions of the major subdivisions within the *Chloroflexi* supergroup as well as ultrafast bootstrap values for the deeply branching nodes which separate the groups. (C) UMAP embedding of a counts matrix representing KOFAM HMM hits across the dataset with taxonomic groups highlighted. Data was Hellinger normalized prior to projection with UMAP.

Within the phylum *Chloroflexota* are four well sampled, deeply branching clades, some of which contain multiple classes but form cohesive phylogenetic groups: the *Chloroflexia*, *Anaerolinea*, *Ktedonobacteria*, and *Dehalococcoidia.* We present new genomes for all four clades. Additionally, rooting proximal to the *Ktedonobacteria* are two deeply branching, poorly sampled lineages, one of which is composed entirely of genomes obtained from coral and sponge holobiont metagenomes^[21][34]^.

Within the class *Dehalococcoidia* are three major groups. Interestingly, two of these groups lack essentially all of the genes necessary for the synthesis of peptidoglycan (**Fig. 2**). This was previously noted for a few cultivated representatives^[22]^ but has not previously been evaluated through an analysis of all publicly available genomes of this clade. One clade within *Dehalococcoidia* includes *Dehaloccoides mccartyi,* which has an S-layer like protein cell wall instead of a wall containing peptidoglycan^[23]^. Another group within *Dehalococcoidia* is the *SAR202* group, primarily comprised of representatives from marine environments^[24][25]^.

**Figure 2.**
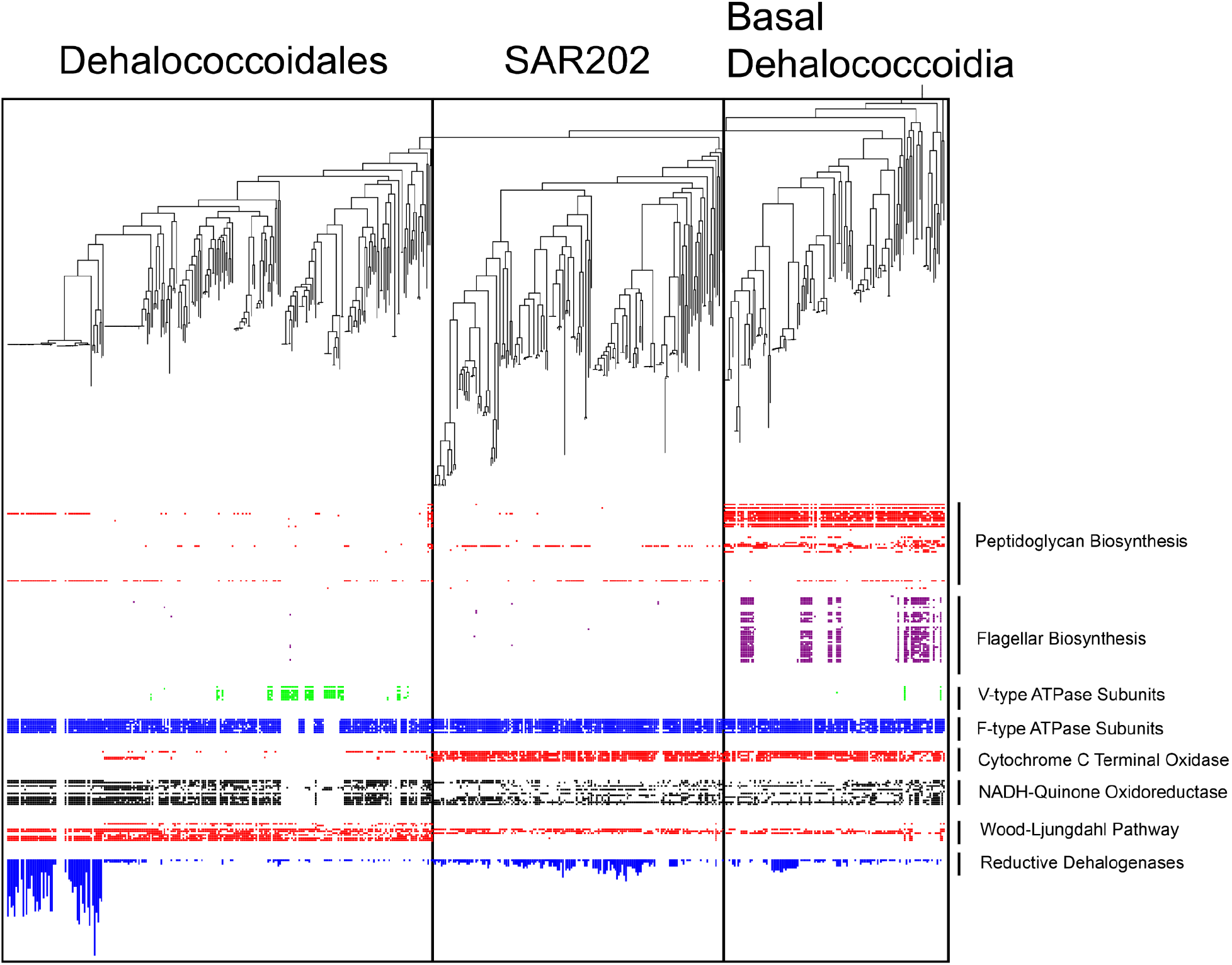
Subsection of the ribosomal phylogeny displayed in figure 1 focusing on the *Dehalococcoidia*, showing the three major functional subdivisions. The Basal Dehalococcoidia lineages are distinguished primarily by their intact peptidoglycan biosynthesis pathways and widely distributed flagellar biosynthesis capacities; the SAR202 lineages lack peptidoglycan biosynthesis capacity but retain many genes required for aerobic metabolism; and the Dehalococcoidales lineages lack markers for aerobic metabolism and contain representatives with high copy numbers of reductive dehalogenases such as the genus *Dehalococcoides* and its close relatives.

Some genomes basal to the *Dehalococcoidia* clade, as well as genomes within the more distal *Dehalococcoidales* clade, contain genes which code for the Wood-Ljungdahl pathway as previously reported^[26]^. The SAR202 clade and the rest of the *Chloroflexi* supergroup lack the genes necessary for the full Wood-Ljungdahl pathway.

### Rubisco

———Ribulose 1,5-bisphosphate carboxylase/oxygenase is widespread in phototrophic and non-phototrophic members of the *Chloroflexi* supergroup. We identified no cases of form II or form II/III Rubisco in the supergroup. All Rubisco large chain (*rbcL*) genes from members of the *Chloroflexi* supergroup are phylogenetically related to form I Rubisco, and form I *rbcL* sequences are found in genomes belonging to all clades analyzed in this study, including *ca. Dormibacteraeota* and *ca. Eulabeiota*. However, one sequence from a groundwater-associated *Eulabeiota* genome and one from a sediment-associated *Eulabeiota* genome occur without *rbcS*, the small subunit canonically found alongside the large chain. These two sequences indicate a clade that is basal to recently reported divergent *Anaerolinea* form 1’ Rubiscos that also lack a small subunit^[13]^. No *rbcS* sequences were found anywhere in the *Eulabeiota* genomes containing these novel *rbcL* sequences. The *Eulabeiota rbcL* sequences display low-level homology to both form III and form I rubisco sequences, at approximately 50% identity to form III representatives and 48% identity to form I representatives by blastP, and cluster with form I representatives in the phylogeny. However, unlike form III rubisco, the Anaerolinea form 1’ Rubiscos and both *Eulabeiota rbcL* genes are encoded adjacent to phosphoribulokinase (*prkB*) and *cbbR*, a transcriptional regulator of the rubisco operon (**Fig. 3**).

**Fig. 3:**
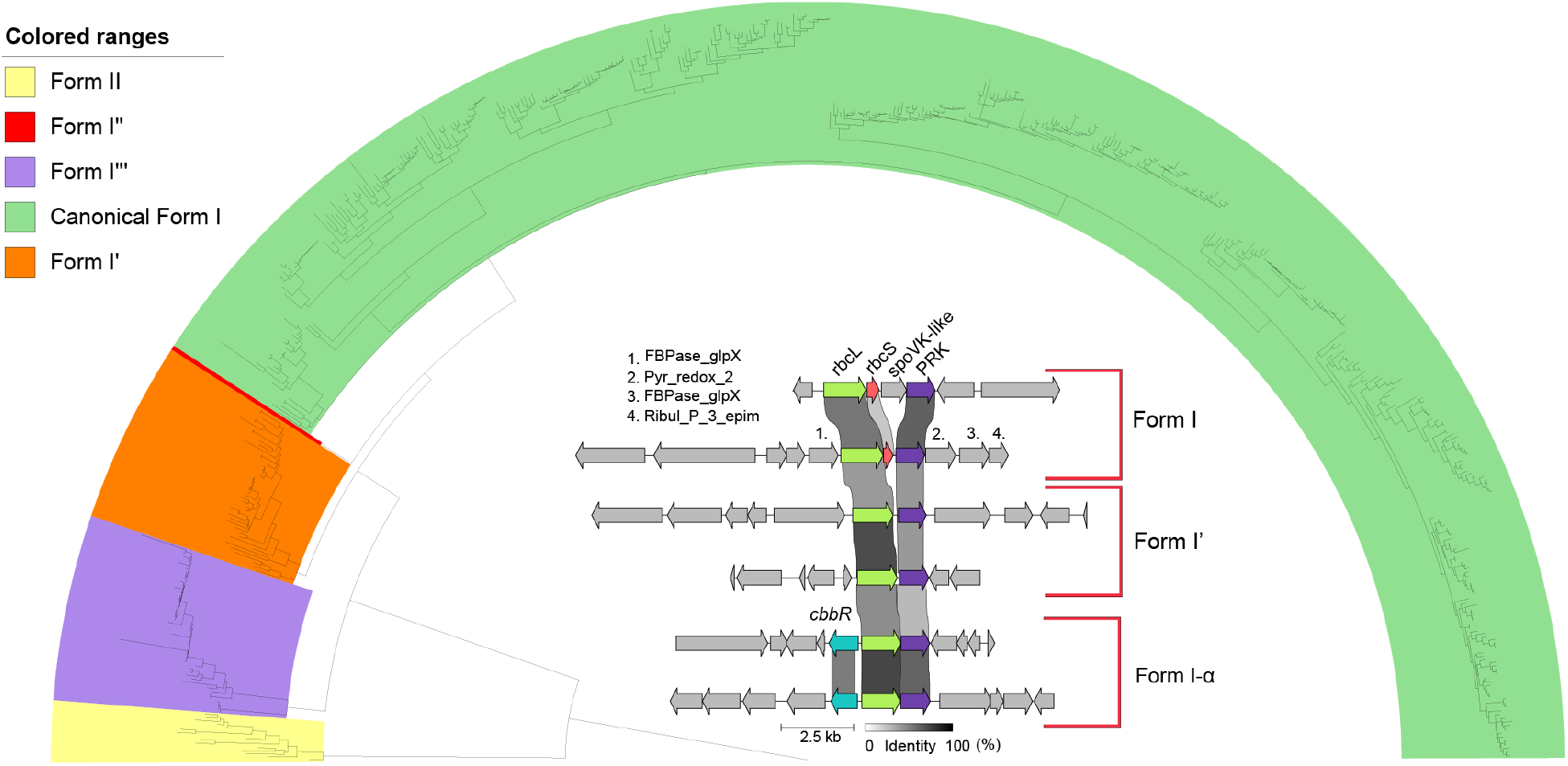
Phylogenetic tree of *rbcL* forms I, I’, I-⍺, with form II used as an outgroup. Interior shows genomic context diagrams for gene clusters containing *rbcL* sequences corresponding to forms I, I’, and 1-⍺, highlighting the presence of PRK in all clusters and lack of *rbcS* in clusters containing *rbcL* sequences of forms I’ and I-⍺.

To further investigate the diversity and environmental distribution of Rubisco with sequences related to those from *Eulabeiota*, we searched a large dataset of unbinned metagenomic data and found examples in datasets from soils, a salt pond and groundwater. (**Fig. 3**). Phylogenetic analysis using a dataset that included these unbinned *rbcL* sequences, the two *Eulabeiota* sequences and sequences form 1’ Rubiscos establish that they form a clade basal to both form 1’ and form I and clearly separate from form III Rubisco. (**Fig. 3a**). We refer to this new clade of Rubisco as form I-*α*. These genomes encode for a partial CBB pathway, and both PRK and PGK are present in these genomes. Given clear phylogenetic affiliation with form I Rubisco, the co-occurrence of the binned and some unbinned sequences with *prkB* and *cbbR*, recent biochemical evidence that form I’ fix CO_2_ despite the absence of *rbcS*, and the presence of PRK and PGK enzymes in the genomes where form 1-*α* is detected, we conclude that the *Eulabeiota* form 1-*α* clade are involved in CO_2_ fixation via the CBB pathway. Additionally, analysis of unbinned *rbcL* sequences revealed a putative clade proximal to form I’ and form I, which we have designated form I’’.

### Photosynthesis

The majority of known phototrophic organisms within the phylum *Chloroflexota* are within the class *Chloroflexia*, and use type II photosystem reaction centers along with bacteriochlorophyll to perform non-oxygenic photosynthesis^[11]^. *Chloroflexota* genomes containing type I photosystem reaction centers have been recently reported^[10]^, and we observed these genes in the genomes deposited to NCBI from this study, but no additional type I photosystem reaction center hits were observed in other genomes from the *Chloroflexi* supergroup. Here we present 6 newly reported genomes from the *Anaerolinea* clade with type II photosystem reaction centers (*pufLM*), and identify 7 *Anaerolinea* genomes with *pufLM* from public data. These photosystem reaction centers show phylogenetic proximity to sequences from the clade *Chloroflexia*, yet separate into clades distinct from sequences of class *Anaerolinea*. *Chloroflexia* and *Anaerolinea* genomes contain *pufLM* fusion genes, as reported previously^[11]^ (**Fig. 4**). Alongside the type II photosystem reaction centers, these genomes contain homologs to photosystem reaction center-associated cytochromes and transcription factors. Additionally, they contain light harvesting proteins associated with the bacteriochlorophyll binding proteins of the chlorosome found in photoautotrophic *Chloroflexia*^[27]^, suggesting that these organisms are capable of performing photosynthesis. These genomes also have at least partial bacteriochlorophyll biosynthesis pathways.

**Figure 4.**
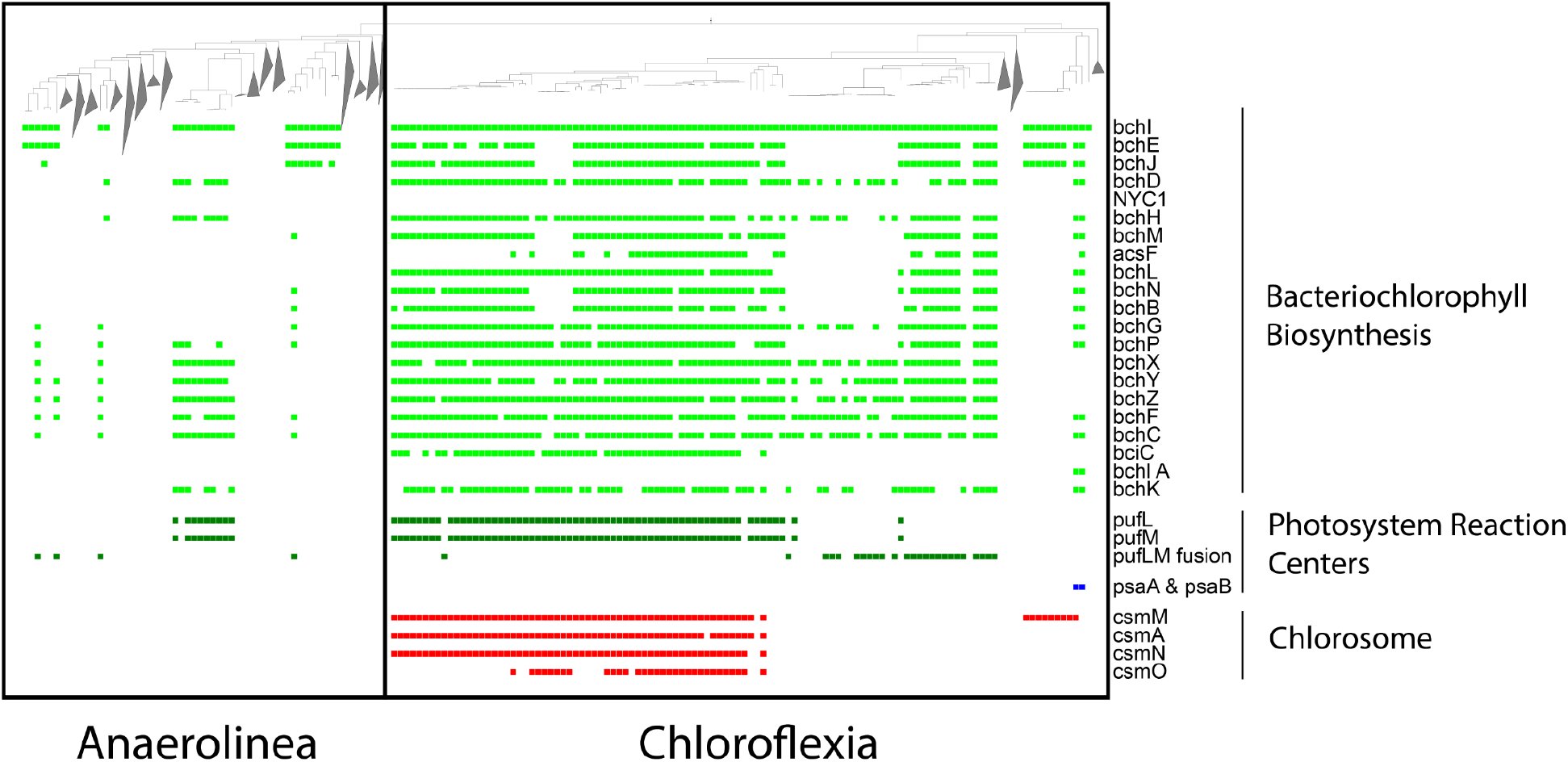
Subsection of the ribosomal phylogeny displayed in Figure 1 highlighting the groups *Anaerolinea* and *Chloroflexia*, which contain phototrophic representatives. Decorations indicate bacteriochlorophyll biosynthesis pathways (light green), photosystem II reaction centers (dark green), photosystem I reaction centers (dark blue), and chlorosome proteins (red).

### Carbon compound utilization

Members of the *Chloroflexi* supergroup vary in their predicted capacities to process carbohydrate compounds. For example, *Anaerolinea* genomes have, on average, nearly 14 times more predicted glycosyltransferases per genome than those of *Dehalococcoidia* (**Fig. 5**). *Anaerolinea* and *Chloroflexia* are enriched in carbohydrate metabolism genes of all CAZy (Carbohydrate-Active enZyme) classes compared to other phyletic groups. Numerous *Dehalococcoidia*, including members of the SAR202 group, and *ca. Dormibacteraeota* have genomes that encode for very few carbohydrate metabolizing enzymes.

**Figure 5.**
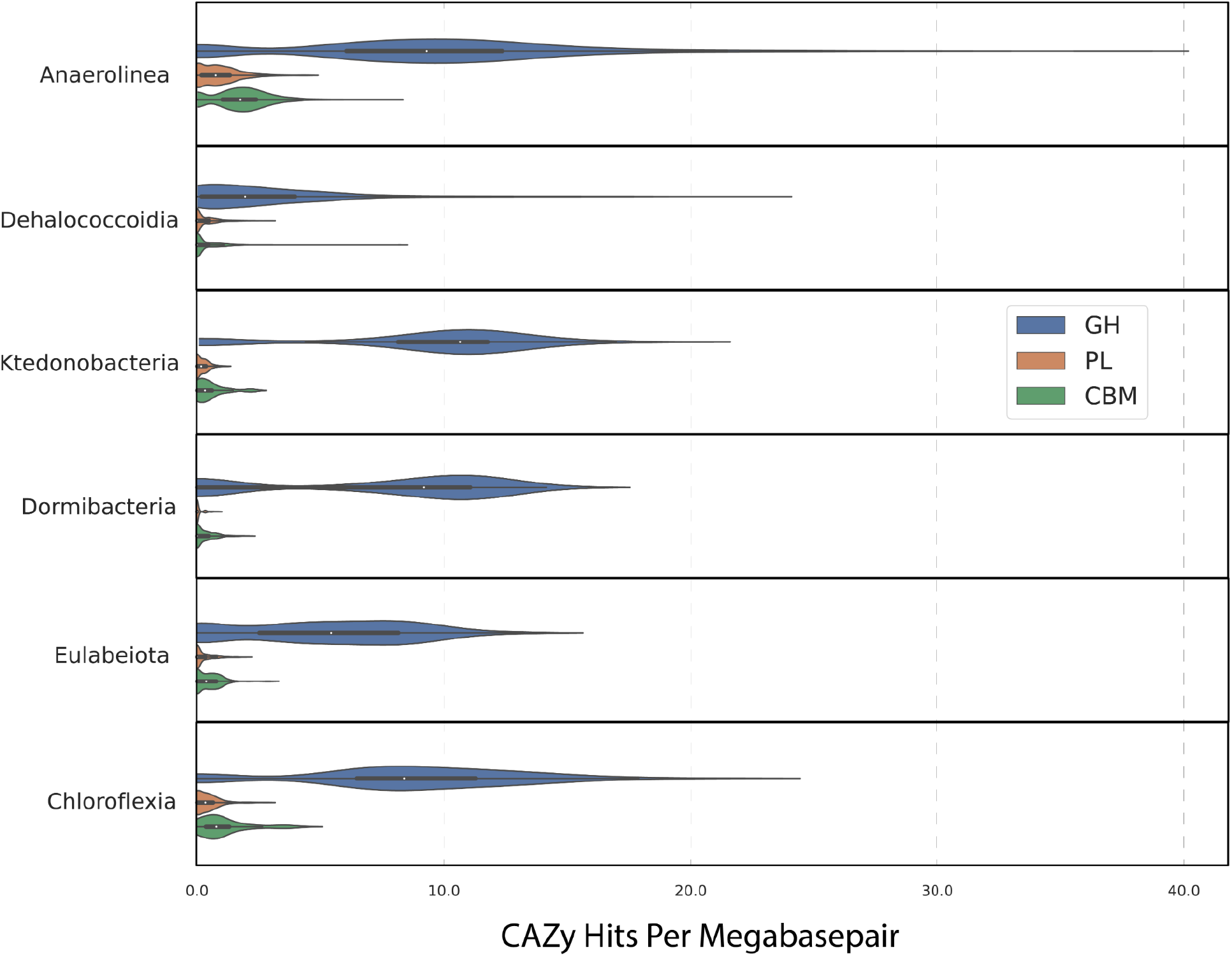
Distribution of CAZy copy number per genome normalized to genome length in megabasepairs for each major clade in the *Chloroflexi* supergroup. CAZy subtypes shown are Glycoside Hydrolase (GH), Polysaccharide Lyase (PL), and Carbohydrate-Binding Modules (CBM).

### Hydrogenases

We investigated the distribution of capacities related to hydrogen metabolism across the Chloroflexi supergroup. To enable predictions, hydrogenases were classified into types using phylogenetic analyses based on the sequences of the large subunit. Detailed information on the hydrogenases observed in genomes from the *Chloroflexi* supergroup is available in **Supplementary table S3** and in **Supplementary Text S4**. Hydrogenases belonging to FeFe group C, FeFe group B, and energy-converting hydrogenase-related (Ehr) complexes are reported here for the first time in the *Chloroflexi* supergroup.

Overall, we find that hydrogenases are abundant, especially in *Anaerolinea* and *Dehalococcoidia*, which are typically found in environments that are anaerobic or periodically anaerobic. The function of these hydrogenases can be H_2_ consumption, utilization, or both. In contrast, the *Eulabeiota* and *Ktedonobacteria* have relatively few hydrogenase hits per genome. The hydrogenases observed within the *Ktedonobacteria* generally belong to NiFe group 1h, which have been observed to oxidize H_2_ at atmospheric concentrations in the presence of O_2_ ^[30]^.

A recently described group of NiFe hydrogenases, group 1l, are found exclusively in the Eulabeiota. These have been hypothesized to provide electrons to Rubisco and support carbon fixation [13]. Consistent with this, the majority of genomes containing group 1l NiFe hydrogenases also contain form I Rubisco (with the small and large subunits), although the hydrogenase and rbc gene clusters are not co-localized in these genomes.

The most numerically abundant hydrogenase subtypes within the *Chloroflexi* supergroup are NiFe group 3, including a novel NiFe group 3c hydrogenase implicated in electron bifurcation, and group 3d NiFe hydrogenases, which usually couple reversible H_2_ oxidation to NAD^+^ reduction (**Fig. 6**). In several *Anaerolinea* and *Dehalococcoidia* genomes, NiFe group 3c hydrogenase was preceded by genes encoding a heterodisulfide reductase (HdrABC), and in some cases electron transfer flavoprotein (ETF) complex. A complex between HdrABC and hydrogenase has been implicated in flavin-based electron bifurcation in Archaea, though it requires a third protein partner ^[28][29]^. An association between group 3c NiFe hydrogenase, EtfAB, and an heterodisulfide reductase (HdrA2B2C2D) has the potential to perform electron bifurcation (see **Supplementary Text S4** for details).

**Figure 6.**
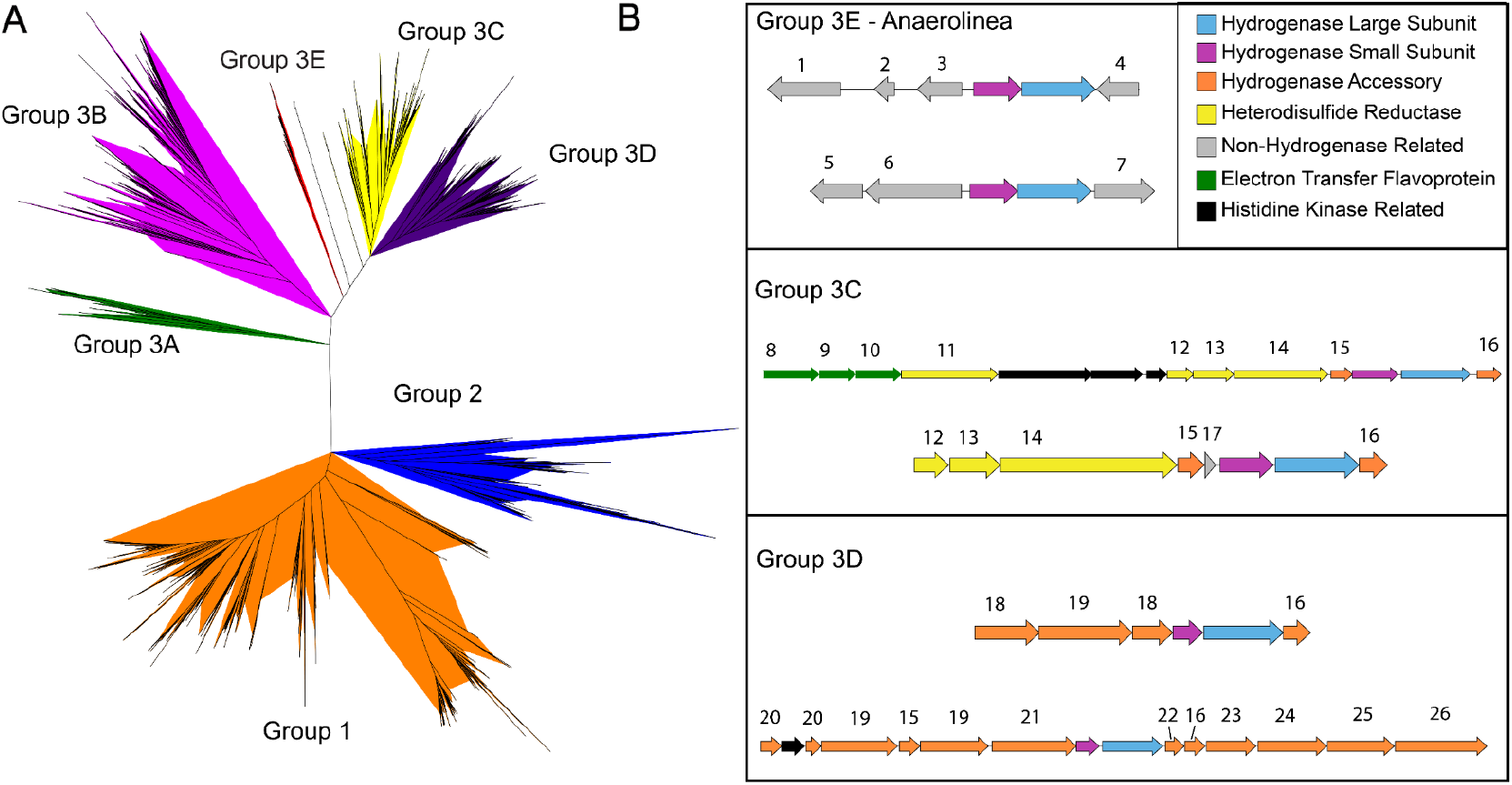
**(A)** Phylogeny of NiFe hydrogenase large subunit proteins from groups 1, 2, and 3, containing sequences from the Chloroflexi supergroup alongside references, and showing the relative placement of the newly proposed group 3E. **(B)** Genomic context diagrams for groups 3E, 3C, and 3D showing the diversity in function between the NiFe hydrogenase subtypes in the Chloroflexi supergroup. Group 3E was not found to regularly co-occur with accessory proteins, whereas group 3C often occurred proximal to heterodisulfide reductase genes (hdrA2B2C2), and group 3D often co-occurred with numerous accessory and electron transfer subunit genes. Numbered genes are annotated as follows: 1. Serpin B, 2. FGE-Sulfatase, 3. Myo-inositol-1 monophosphatase, 4. ADP-ribosylglycohydrolase, 5. UDP-Glucose-4-epimerase, 5. Ubiquinone biosynthesis protein, 6. Peptidase M23, 7. ETF-QO (electron acceptor subunit), 8. ETF Beta subunit, 9. ETF Alpha subunit, 10. Heterodisulfide reductase subunit D, 11. Heterodisulfide reductase subunit C2, 12. Heterodisulfide reductase subunit B2, 13. Heterodisulfide reductase subunit A2, 14. Hydrogenase Fe-S subunit, 15. HycI protease, 16. PIN domain protein, 17. Bidirectional [NiFe] hydrogenase diaphorase subunit, 18. nqoF-like, 19. NADP-reducing hydrogenase subunit hndB, 20. nqoG, 21. CBS domain-containing protein, 22. Formate dehydrogenase Fe-S binding subunit, 23. Formate dehydrogenase subunit alpha, 24. LysM motif-containing metalloendopeptidase, 25. coxL family molybdopterin aldehyde dehydrogenase.

Unique to the *Anaerolinea* is a divergent clade of NiFe group 3 hydrogenases for which the large subunit is phylogenetically proximal to groups 3c and 3d. For this group, we propose the group name 3e. This hydrogenase is accompanied by a small subunit but lacks both identified accessory proteins and electron transfer subunits, and may interact with unknown partners.

In all groups, we identified hydrogenases that likely support H_2_-oxidation-based energy generation. Other hydrogenases, especially those associated with the cell membrane (NiFe type 4), are likely involved in proton translocation and were found across the Chloroflexi supergroup, with the exception of the *Dormibacteraeota.* (**Supplementary figure S3, Supplementary Text S4**)

### Multiheme Cytochromes

Multiheme cytochrome proteins, defined as having ≥ 4 heme-binding motifs, are especially abundant in Anaerolinea. Of all proteins with ≥ 20 CxxCH heme-binding motifs in the Chloroflexi supergroup, the majority (88 of 124) occur in genomes from the *Anaerolinea*. Large multiheme cytochromes with ≥20 heme-binding motifs are understood to participate in redox reactions, for example iron and manganese reduction reactions^[31]^, and are hypothesized to play a role in extracellular respiration in iron reducing bacteria^[32]^. Many of the large multiheme cytochromes contain transmembrane domains and demonstrate homology to models trained on multiheme cytochrome proteins from the genus *Geobacter* (GSu_C4xC_C2xCH), noted for its ability to participate in extracellular metal ion transformations. This suggests that Anaerolinea use multiheme cytochromes to deliver electrons to an extracellular terminal electron acceptor.

### Nitrogen cycling

Within the *Chloroflexi* supergroup, genes encoding for nitrogen transformation pathways are common but unevenly distributed phylogenetically (**Fig. 7**). Several nitrogen transformation capacities were not detected in the *Chloroflexi* supergroup, including ammonia oxidation (*amoAB*), nitrite oxidation by *nasAB*, and N_2_O reduction by *norBC*.

**Figure 7.**
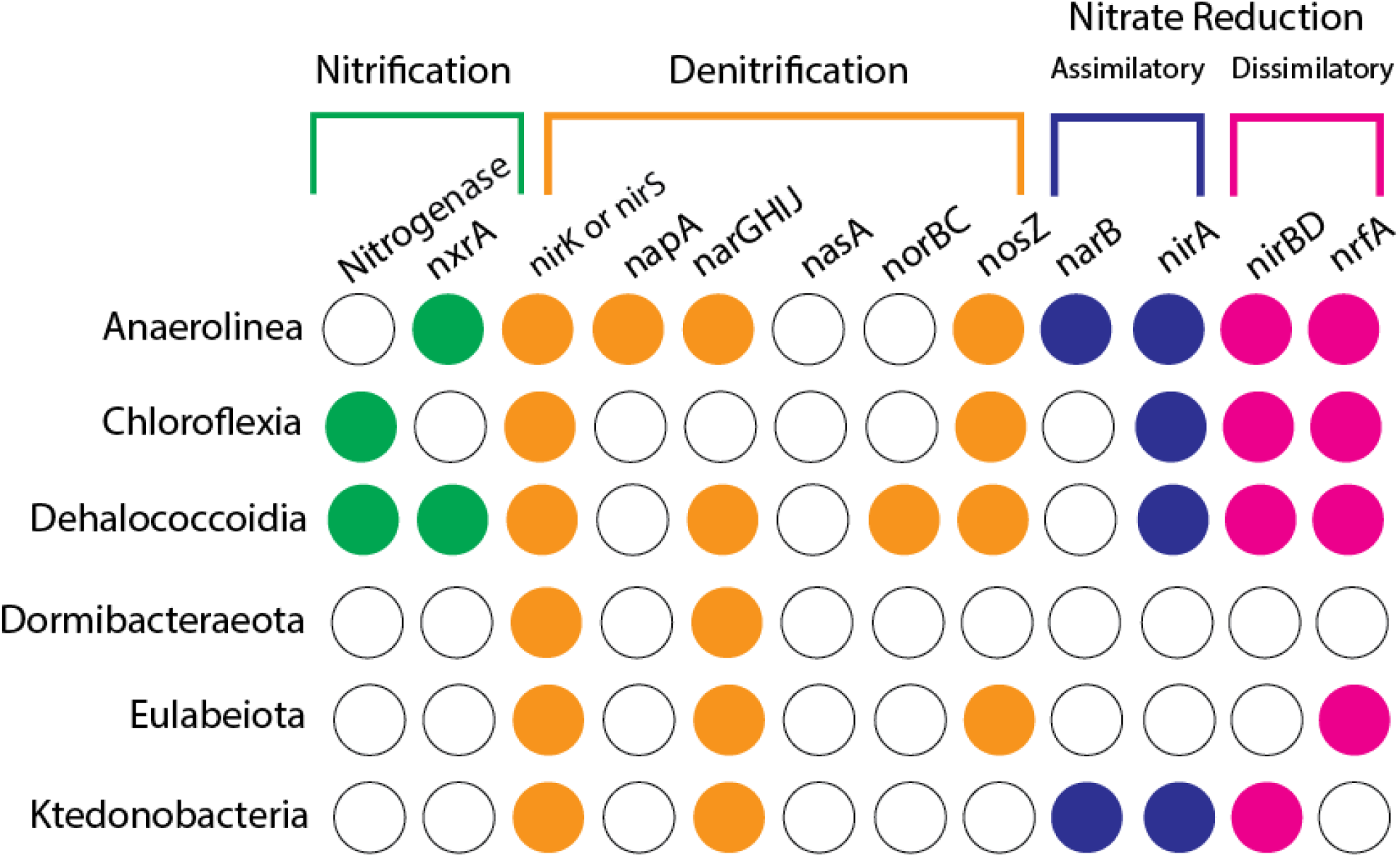
Colored circles indicate the presence or absence of important nitrogen cycling genes in major subdivisions of the *Chloroflexi* supergroup.

Reduction of nitrate and nitrite are the most commonly observed nitrogen transformation capacities in the *Chloroflexi* supergroup; all studied clades contained representatives capable of reducing nitrite. Only *ca. Dormibacteraeota* lack the genomic capacity to perform assimilatory or dissimilatory nitrate reduction. There are many *Anaerolinea* and *Dehalococcoidia* that can reduce nitrogen-containing compounds, and via many mechanisms. In contrast, *Dormibacteraeota* encode relatively few genes for nitrogen compound transformations, and a low diversity of pathways.

Some *Anaerolinea* genomes contain multiple genes for reduction of NO_3_^−^ to NO_2_^−^. They contain the *narGHIJ* nitrate reductase complex, periplasmic nitrate reductase (*napAB*), which is part of the dissimilatory pathway, and ferredoxin nitrate reductase (*narB*), which is involved in assimilatory nitrate reduction. The *narGHIJ* system is also present in genomes from the candidate phyla *Dormibacteraeota* and *Eulabeiota,* but was not detected in class *Chloroflexia* or *Ktedonobacteria.*

Reduction of NO_2_^−^ to NH_3_ through the *nrfAH* denitrification system is very common within the *Anaerolinea*. This pathway was identified even where genes for nitrate reduction were not identified elsewhere in the genome, suggesting that these organisms rely on nitrite produced by coexisting community members.

Nitrite reduction via *nirA*, associated with the assimilatory pathway, is present in all clades of the phylum *Chloroflexota* but absent from the candidate phyla *Dormibacteraeota* and *Eulabeiota*. The capacity for NO_2_^−^ reduction to NO via *nirK* or *nirS* is common across the supergroup. The capacity to reduce NO to N_2_O via *norBC* is only observed in the *Dehalococcoidia*. Further reduction of N_2_O to N_2_ via *nosZ* is common across the Chloroflexi supergroup, but absent from the *Ktedonobacteria* and *ca. Dormibacteraeota*.

Nitrogen fixation is present in the classes *Dehalococcoidia* and *Chloroflexia*. Notably, there are multiple phylogenetically distinct forms of *nifH* (**Supplementary Fig. S4**). The *Dehalococcoidia* containing *nifH* are distal to those found in *Chloroflexia* and cluster with group II *nifH* sequences, whereas nitrogenases from class *Chloroflexia* form a clade separate from previously defined subtypes basal to form I *nifH*. All of the *nifH* sequences observed in genomes also containing *nifA* and *nifB* are further implicated in nitrogen fixation by phylogenetic proximity to biochemically verified sequences.

### The Sulfur Cycle

Microorganisms from the *Chloroflexi* supergroup are well known to participate in biogeochemical sulfur cycling^[32]^, but the distribution of sulfur cycle genes throughout the supergroup has not been reported. Our analysis indicates that the most common genes are implicated in assimilatory sulfate reduction. The *sat* sulfate adenylyltransferase system, which can be involved in both assimilatory and dissimilatory steps, is common throughout the *Chloroflexi* supergroup. An alternative gene for this reaction, sulfate reductase, PAPSS/K13811, is found primarily in genomes of *ca. Dormibacteraeota,* possibly as part of the assimilatory sulfate reduction pathway. The assimilatory reduction of SO_3_ ^2-^ to H_2_S via the *cysIJ* and *sir* sulfite reductase systems seems to be particularly common among the *Eulabeiota* and *Ktedonobacteria*, but absent from *Dormibacteraeota*.

The reduction of adenylylsulfate to SO_2_ ^−^ via *aprAB* in the dissimilatory sulfate reduction pathway is rare and sparsely distributed in the *Chloroflexi* supergroup. The *dsrAB* genes, which can both reduce sulfite and oxidize sulfide, are rare. Sulfur oxygenase-reductase were identified in two *Anaerolinea* genomes, but this gene is absent from the rest of the supergroup. Thiosulfate oxidation via the *sox* complex was not predicted for any bacteria of the Chloroflexi supergroup.

Dimethyl sulfoxide (DMSO) reductase genes of the *dmsABC* family were detected in all lineages of the *Chloroflexi* supergroup with the exception of the *Dormibacteraeota*, although genes that would indicate metabolism of the DMSO breakdown product dimethyl sulfide (DMS), such as dimethyl-sulfide monooxygenase (*dmoA*) and dimethyl sulfide dehydrogenase (*ddhABC*), were not detected in this dataset. Undetected in the dataset were genes implicated in the breakdown of dimethylpropiothetin (DMSP, a compatible solute) to form DMS. The only gene present in the pathway that converts DMS ultimately to sulfate (e.g. *sox* and *sor*) is methanethiol oxidase, which produces sulfide from methylmercaptan, and is particularly abundant in the *Dormibacteraeota* and *Ktedonobacteria*.

### CRISPR-Cas Systems

The abundance of CRISPR-Cas phage defense systems within the *Chloroflexi* supergroup is highly dependent upon the environment of origin and organism taxonomy (**Supplementary Figure S5**). The candidate phyla *Dormibacteraeota* and *Eulabeiota* have a strikingly low average number of detected CRISPR arrays relative to other groups, although intact cas protein cassettes are detected in genomes from these candidate phyla which otherwise lack detectable CRISPR arrays.

## Discussion

Taxonomic databases disagree on the number of subdivisions within the Chloroflexi (alternatively called the *Chloroflexota*). For simplicity, we used the long-established phylogenetically cohesive class-level subdivisions *Anaerolinea*, *Dehalococcoidia*, *Ktedonobacteria*, and *Chloroflexia* as the foundation for our further taxonomic analyses. Our ribosomal phylogeny clearly supports the existence of these four main groups. A poorly sampled lineage which lies proximal to the *Ktedonobacteria* (**Fig. 1**) is comprised entirely of endosymbionts of sponges and corals ^[21][34]^.

External to the *Chloroflexota* lie the candidate phylum *Dormibacteraeota (*previously Candidate Division AD3) and another distinct lineage that includes groups previously named *Limnocylindria* and *Edaphomicrobia*, additionally demarcated by GTDB as class *Ellin6529* within the phylum *Chloroflexota*. The *Limnocylindria* and *Edaphomicrobia were* recently proposed classes within the Chloroflexota. Our analyses indicate that the *Limnocylindria* and *Edaphomicrobia* are closely related, part of a single phylum-level lineage, and that they place outside of the Chloroflexota. Our newly reported genomes clarify their phylogeny and substantially expand the breadth of the lineage that includes the newly proposed candidate phylum *ca. Eulabeiota*. Together, the Chloroflexota, *Eulabeiota* and *Dormibacteraeota comprise* the Chloroflexi supergroup, which is part of the larger *Terrabacteria* group.

### Eulabeiota

The *Eulabeiota* were first named Ellin6529 after an isolate was obtained from agricultural soil near Ellinbank, Victoria, Australia^[36]^. Unfortunately, the isolate was lost. However, a full-length 16S rRNA sequence was obtained and used to study the abundance of this lineage across habitats. Although the majority of *Eulabeiota* sequences we studied came from soils, often from cold or perennially dry environments, they also appear in freshwater environments, such as partially melted permafrost^[5]^, lakes^[7]^, and groundwater^[38][39][6]^ [Christine’s paper, BJP, relevant Rifle publications]. Notably, genomes from the *Eulabeiota* have been obtained from marine oil seeps^[81]^ and one from a hydrothermal vent^[37]^.

The *Eulabeiota* lack outer membrane synthesis genes and contain similar peptidoglycan biosynthesis machinery to their sister phyla *Chloroflexota* and *Dormibacteraeota* as well as more distant neighbors such as the *Actinobacteria* and *Armatimonadetes*, suggesting that the *Eulabeiota* are monoderm organisms.

Nearly all *Eulabeiota* genomes encode at least one type of cytochrome *c* terminal oxidase, indicating that they are at least facultatively aerobic. However, some genomes lack terminal oxidases, and have markers for anaerobic metabolism such as anaerobic CO dehydrogenase (*cooS*), suggesting that some *Eulabeiota* have adapted to strictly anaerobic environments.

Some *Eulabeiota* genomes encode the capacity for CO_2_ fixation via the CBB pathway using Rubisco (*rbcL*). Interestingly, phylogenetic analysis separates the Rubisco genes present in the *Eulabeiota* into multiple clades, one of which, designated form I-⍺, lacks the small subunit *rbcS* (**Fig. 3**). Both genomes with the I-⍺ rbcL were obtained from aquifer water and sediment from Rifle, Colorado^[6]^, in an environment with low dissolved oxygen content. This Rubisco clade is more deeply branching than the other recently reported single subunit form I’ Rubisco, which was shown biochemically to function as an oxygen-sensitive carboxylase enzyme in the absence of rbcS^[13]^. We infer that I-⍺ rbcL is similar in structure and function to the precursor of modern Rubisco found in cyanobacteria, algae, and plants. Consistent with the suggestion for form I’ Rubisco, we infer that the Eulabeiota form I-⍺ evolved to function in an anaerobic world and that this precursor gave rise to the two subunit form I complex that was widely laterally transferred across the tree of life after the rise of O_2_ in the atmosphere.

### Dormibacteraeota

The recently characterized candidate phylum *Dormibacteraeota* has thus far been found exclusively in soil and permafrost metagenomes, and has no cultured representatives. This clade seems to be comprised of at least facultatively aerobic monoderm organisms, as evidenced by a widespread distribution of aerobic CO dehydrogenase gene cassettes as well as cytochrome C terminal oxidase genes throughout the phylum. Hydrogen metabolism is also abundant, with *Dormibacteraeota* genomes containing type 1 and type 3 NiFe hydrogenases, although type 1 and type 3 NiFe hydrogenase subtypes are not observed together in a single genome from this clade (**Supplementary Table S3**).

### Dehalococcoidia

The *Dehalococcoidia* contains three major phylogenetic subdivisions that exhibit different sets of metabolic pathways and have characteristic environmental distribution patterns (**Fig. 2**). We will refer to these three subdivisions as the basal clade, the *SAR202*, and the *Dehalococcoides* clade (**Fig. 2**).

Notable is the absence of a functional peptidoglycan biosynthesis pathway in a substantial portion of the*SAR202* and *Dehalococcoides* clades. However, genomes in the basal clade often have peptidoglycan biosynthesis pathways. Previous studies^[8]^ have highlighted a lack of peptidoglycan in *Dehalococcoides mccartyi*, the type species of the *Dehalococcoidia*, although microscopy reveals a cell wall-like structure perhaps similar in function to those with S-layers found in other bacteria and archaea.

Some genomes within the *Dehalococcoides* lack F-type ATPases, instead containing only a V-type ATPase, as has been previously reported^[26]^. The genomes that contain V-type and lack F-type ATPases, along with a small subset of other *Dehalococcoides* genomes, lack genes for the NADH-quinone oxidoreductase complex, indicating the use of substrate-level phosphorylation to generate ATP.

The functional division within the class *Dehalococcoidia* also correlates with the environments of origin. The *SAR202* clade are primarily from marine or saline environments, whereas genomes from the *Dehalococcoides* and basal clades are generally from terrestrial sources such as groundwater, soil, and freshwater, although there are exceptions in each case.

### Anaerolinea

The most well-represented clade within the dataset is the *Anaerolinea*. They have extensive capacity for nitrogen metabolism, some are likely photosynthetic and some are capable of assimilatory and dissimilatory sulfate reduction. Many are predicted to be autotrophs, using both Form I and Form 1’ Rubisco in the CBB pathway, and Form 1’ is exclusively found in genomes which fall in this clade. Many *Anaerolinea* genomes encode numerous multiheme cytochromes, some of which have >20 heme-binding motifs per protein. These may be involved in extracellular electron transfer reactions, including metal reduction and oxidation^[88][89]^.

Photosystem II reaction centers phylogenetically distinct from those found in the *Chloroflexia* are observed in genomes from the *Anaerolinea*. These genomes contain partial bacteriochlorophyll synthesis pathways but include crucial genes such as *bchP*, which catalyzes the last reaction in the biosynthesis of bacteriochlorophyll *c*, suggesting that although the full bacteriochlorophyll synthesis pathway is not detected in these genomes that the synthesis of bacteriochlorophyll does occur, either using divergent bacteriochlorophyll synthesis genes or by obtaining precursor compounds from other bacteria in their communities. These putatively photosynthetic *Anaerolinea* are primarily found in hot spring environments, although three such genomes were obtained from stromatolite metagenomes^[12][40]^.

The *Anaerolinea* contain a number of unique subtypes of NiFe hydrogenases, including the phylogenetically divergent group 3e, which is unique to the *Anaerolinea*, and a variant of group 1f which lacks a cytochrome subunit but associates with a *nrfD*-like molybdopterin subunit. They are the only clade within the *Chloroflexi* supergroup to contain FeFe hydrogenases of type B. *Anaerolinea* genomes also contain hydrogenases which are proximal to heterodisulfide reductase complexes as well as electron transfer flavoprotein subunits, potentially implicating these organisms in electron bifurcation. The diversity in the number of observed hydrogenase subtypes in the *Anaerolinea*, as well as the unique hydrogenase subtype of group 3e observed exclusively therein, points to the importance of hydrogen-based metabolism for members of this lineage, and suggests that members of this clade are well-adapted to at least periodically anaerobic environments.

*Anaerolinea* have the most varied environmental distribution of any clade in this study, with collection temperatures ranging from hot springs and hydrothermal vents^[41]^ to permafrost ^[5]^ and the human oral microbiome (BioProject PRJNA282954). They are common in soils and occur in activated sludge from wastewater treatment plants^[42]^, where they contribute to sludge flocculation. Their notably extensive metabolic diversity may in part reflect their adaptation to very diverse environments, and the associated varied energy resources.

### Chloroflexia

The *Chloroflexia* group is well-studied, and contains the class *Chloroflexia*, the type class of the *Chloroflexota*. The group contains the majority of the phototrophic organisms within the phylum, and is unique among the clades of the *Chloroflexi* supergroup in that several are capable of performing CO_2_ fixation via the 3-hydroxypropionate bicycle, as has been previously observed^[43]^. Genomes from this group, especially those most closely related to *Chloroflexus aggregans*, the photoautotrophic type species of the phylum, tend to be observed most often in hot springs or freshwater sources. More divergent lineages within the group basal to the *Chloroflexia* are more often found in soil, and lack genes coding for anoxygenic photosynthetic reaction centers as well as the complete 3-hydroxypropionate bicycle, suggesting they occupy different metabolic niches despite their phylogenetic proximity to typical *Chloroflexia*.

### Ktedonobacteria

The *Ktedonobacteria* group is comprised of two phylogenetically distinct subdivisions, one of which is comprised entirely of genomes obtained from soil samples and includes the type genus *Ktedonobacter*, and another comprised of genomes primarily from stratified freshwater lakes and ponds^[44]^ and groundwater sources^[38]^. *Ktedonobacteria* genomes are of particular research interest because of their size and the large number of secondary metabolite gene clusters. This extends prior findings^[45][46]^ which show that substantial inventories of secondary metabolite gene clusters in genomes obtained from soil. Products of this type of metabolism may be implicated in interaction among organisms or between organisms and their environment.

*Ktedonobacteria* contain fewer hydrogenase subtypes than other clades, and the majority of the observed subtypes in this clade belong to the O_2_-tolerant type 1h^[30]^. Hydrogen oxidation is likely important in anaerobic (e.g., groundwater) or seasonally anaerobic soils where fermentation generates H_2_. Members of this group also contain form I *rbcL* sequences which coincide with *rbcS* and phosphoribulokinase genes in the same genome, strongly suggesting that these organisms are capable of carboxydotrophy.

## Conclusion

The *Chloroflexi* supergroup is comprised of bacteria from across diverse environments, and is largely understood by way of genome-resolved metagenomics. Our study provides the most detailed ribosomal phylogeny of the *Chloroflexi* supergroup to date, which allowed for an investigation of the distribution of important functional genes in clades throughout the supergroup and revealed functional differences between and within class- and phylum-level lineages within the supergroup. Many genomes belonging to the *Chloroflexi* supergroup contain novel genomic features, which are specific to particular clades within the supergroup and contain phylogenetically novel representatives of well-studied protein families such as Rubisco and NiFe hydrogenases. Our work highlights the diversity and ubiquity of hydrogen-dependent metabolism in the *Chloroflexi* supergroup and reveals phylogenetically novel clades of putative hydrogenases of type 3e which have thus far only been observed in genomes belonging to the supergroup. Additionally, we report for the first time the phylogenetic distribution of multiple *Chloroflexi* supergroup-exclusive clades of form I-like Rubisco as first reported in Banda et al.^[13]^, including a new form of putative Rubisco designated form I-⍺. Biochemical investigation of novel proteins such as form I-⍺ Rubisco and group 3e NiFe hydrogenase is crucial to understand the potentially significantly altered catalytic function of these groups relative to biochemically characterized clades. Our results should guide targeted cultivation efforts and aid further investigations into these ubiquitous and understudied organisms.

## Methods

### Database Construction

Genomes were downloaded from three databases: NCBI GenBank^[47]^, PATRIC^[48]^, and ggKbase (https://ggkbase.berkeley.edu/). All database downloads were performed on August 15, 2020. Those downloaded from NCBI GenBank were downloaded using a custom script (attached, supplemental) which utilizes the NCBI Entrez python API, and searched for all genbank genomes with hits to ‘Chloroflexi’. Genomes from BioProjects less than two years old and without associated publications were discarded. Genomes from PATRIC were gathered by searching the keywords ‘Chloroflexi’ and ‘AD3’, the former name for the Dormibacteraeota, on PATRIC and downloading all resulting genomes on Dec. 13, 2019. Genomes from ggKbase were downloaded using only genomes with taxonomic hits to *Dormibacteraeota* (ANG-CHLX), *Eulabeiota* (RIF-CHLX), or *Chloroflexota*. Information on the originating database and additional metadata for each genome can be found in **Supplementary Table S1**.

Genomes from all sources were then dereplicated using dRep^[49]^ using a 100% identity dereplication threshold, additionally removing genomes with greater than 25% checkM^[50]^ contamination and less than 75% checkM completeness.

Genomes were annotated using KOFAMScan^[51]^, applying provided bitscore thresholds. KOFAMscan hits were then counted using a custom python script. Counts were normalized using the Hellinger transformation^[52]^ prior to projection with Uniform Manifold Approximation and Projection (UMAP)^[53]^. Genomes were additionally annotated using USEARCH[84] against Uniprot ^[85]^, Uniref90 and KEGG^[86]^, and 16S rRNA and tRNAs predicted as described in Diamond et al.^[4]^.

All phylogenetic trees included in this paper were visualized using the Interactive Tree of Life (iTOL)^[54]^.

Newly presented genomes in this study were obtained from four projects and processed using the ggKbase annotation and binning pipeline. Sampled locations include: hot springs in Tibet and Yunnan province, China; deep boreholes in Japan (BJP); soil samples taken from the East River watershed in Colorado, United States; and a series of anammox and dechlorination bioreactors.

### Analyses for samples obtained from the Borehole Japan Project

Genomes from the Borehole Japan project were obtained from ~439 L of groundwater samples collected at the Horonobe Underground Research Laboratory and the Mizunami Underground Research Laboratory in Japan, according to methods outlined in Hernsdorf et al. 2017^[39]^ and Matheus Carnevali et al.,2019^[67]^. In brief, genomic DNA was extracted from the biomass gathered on the 0.22 μm GVWP filters using an Extrap Soil DNA Kit Plus version 2 (Nippon Steel and Sumikin EcoTech Corporation, Tsukuba, Japan). Genomic DNA libraries were generated using TruSeq Nano DNA sample Prep Kit (Illumina, San Diego, CA, USA) according to the manufacturer’s instructions, and 150 bp paired-end reads with a 550 bp insert size were sequenced by Hokkaido System Science Co. using Illumina HiSeq 2500. Assembly and binning were performed as reported previously^[39][67]^.

### Sampling, DNA extraction, metagenomic sequencing and analyses for soil samples obtained from the East River, Crested Butte, CO

Genomes were obtained from soil cores sampled from the East River in Crested Butte, CO in 2016 and 2017. Samples were taken from two depth regimes: shallow (between 3-10cm from the soil surface) and deep (between 9-20cm from the soil surface). DNA was extracted from the samples using the Qiagen Powermax Soil DNA extraction kit and submitted to the Joint Genome Institute for sequencing. Soil samples were collected using sterile tools, including a soil core sampler and 7.6 × 15.2 cm plastic corer liners (AMS, Inc), stainless-steel spatulas, and Whirl-pak bags. Samples were immediately stored in coolers for transportation to RMBL, where samples were prepared for archival and transportation to the University of California, Berkeley. Soil cores were broken apart and manually homogenized inside the Whirl-pak bags. Subsamples for chemical analyses, DNA extractions, and long-term archival were obtained inside a biosafety cabinet, kept at − 80 °C, transported in dry ice, and stored at − 80 °C at the University of California, Berkeley.

Genomic DNA was extracted from ~ 10 g of thawed soil using Powermax Soil DNA extraction kit (Qiagen) with some minor modifications as follows. Initial cell lysis by vortexing vigorously was substituted by placing the tubes in a water bath at 65 °C for 30 min and mixing by inversion every 10 min to decrease shearing of the genomic DNA. After adding the high concentration salt solution that allows binding of DNA to the silica membrane column used for removal of chemical contaminants, vacuum was used instead of multiple centrifugation steps. Finally, DNA was eluted from the membrane using 10 mL of the elution buffer (10 mM Tris buffer) instead of 5 mL to ensure full release of the DNA. DNA was precipitated out of solution using 10 mL of a 3-M sodium acetate (pH 5.2) and glycogen (20 mg/mL) solution and 20 mL 100% sterile-filtered ethanol. The mix was incubated overnight at 4 °C, centrifuged at 15,000 × *g* for 30 min at room temperature, and the resulting pellet was washed with chilled 10 mL sterile-filtered 70% ethanol, centrifuged at 15,000 × *g* for 30 min, allowed to air dry in a biosafety cabinet for 15–20 min, and resuspended in 100 μL of the original elution buffer. Genomic DNA yields were between 0.1 and 1.0 μg/μL except for two samples with 0.06 μg/μL. Power Clean Pro DNA clean up kit (Qiagen) was used to purify 10 μg of DNA following manufacturer’s instructions except for any vortexing which was substituted by flickering of the tubes to preserve the integrity of the high-molecular-weight DNA. DNA was resuspended in the elution buffer (10 mM Tris buffer, pH 8) at a final concentration of 10 ng/μL and a total of 0.5 μg of genomic DNA. DNA was quantified using a Qubit double-stranded broad range DNA Assay or the high-sensitivity assay (ThermoFisher Scientific) if necessary. Additionally, the integrity of the genomic DNA was confirmed on agarose gels and the cleanness of the extracts tested by absence of inhibition during PCR.

⸏Clean DNA extracts and co-extracts were submitted for sequencing at the Joint Genome Institute (Walnut Creek, CA), where samples were subjected to a quality control check. Sequencing libraries were prepared in microcentrifuge tubes. One hundred nanograms of genomic DNA was sheared to 600 bp pieces using the Covaris LE220 and size selected with SPRI using AMPureXP beads (Beckman Coulter). The fragments were treated with end repair, A-tailing, and ligation of Illumina-compatible adapters (IDT, Inc) using the KAPA Illumina Library prep kit (KAPA biosystems). Libraries for the rest of the samples were prepared in 96-well plates. Plate-based DNA library preparation for Illumina sequencing was performed on the PerkinElmer Sciclone NGS robotic liquid handling system using Kapa Biosystems library preparation kit. Two hundred nanograms of sample DNA was sheared to 600 bp using a Covaris LE220 focused-ultrasonicator. The sheared DNA fragments were size selected by double-SPRI and then the selected fragments were end-repaired, A-tailed, and ligated with Illumina-compatible sequencing adaptors from IDT containing a unique molecular index barcode for each sample library.

All the libraries were quantified using KAPA Biosystem’s next-generation sequencing library qPCR kit and a Roche LightCycler 480 real-time PCR instrument. The quantified libraries were then multiplexed with other libraries, and the pool of libraries was prepared for sequencing on Illumina HiSeq sequencing platform utilizing a TruSeq paired-end cluster kit, v4, and Illumina’s cBot instrument to generate a clustered flow cell for sequencing. Sequencing of the flow cell was performed on the Illumina HiSeq 2500 sequencer using HiSeq TruSeq SBS sequencing kits, v4, following a 2 × 150 indexed run recipe.

Methods used for metagenome assembly and annotation are described elsewhere [82]. In brief, after quality filtering, reads from individual samples were assembled separately using IDBA-UD v1.1.1 [80] with a minimum k-mer size of 40, a maximum k-mer size of 140, and step size of 20. Only contigs > 1 Kb were kept for further analyses. Reads were mapped to the assemblies using Bowtie2^[55]^ and default settings to estimate coverage.

Annotated metagenomes from both years were uploaded onto ggKbase (https://ggkbase.berkeley.edu), where binning tools based on GC content, coverage, and winning taxonomy ^[38]^ were used for genome binning. These bins and additional bins that were obtained with the automated binners ABAWACA1 (https://github.com/CK7/abawaca), ABAWACA2, MetaBAT^[56]^, Maxbin2^[91]^, and Concoct ^[92]^ were pooled, and DAStool ^[93]^ was used for selection of the best set of bins from each sample as described by Diamond et al.^[4]^, with a completeness threshold applied of >=70% as measured by checkM^[50]^.

### Sampling, DNA extraction, metagenomic sequencing and analyses for dechlorinating and anammox bioreactors

Genomes were obtained from reactors described in Lee et al. 2019^[77]^ and Mao et al. 2020^[78]^. Genomic DNA was extracted from the samples using either the Qiagen (Valencia, CA, USA) DNeasy Blood & Tissue kit or the AllPrep DNA/RNA Mini Kit (Qiagen) according to the manufacturer’s recommendations as outlined in Lee et al. 2020. Metagenomic reads were assembled with IDBA_UD^[80]^ with default parameters. Metagenomic binning was then performed using tetranucleotide frequency ESOMs^[79]^ and the ggkbase manual binning platform.

### Sampling, DNA extraction, metagenomic sequencing and analyses for Tibet and Yunnan hot springs

Hot spring sediment samples were collected in 2016 from Tibet Plateau and Yunnan Province, China. The microbial community and structure in those samples have been reported previously (Chen et al. 2019)^[41]^. Samples were collected from the hot spring pools using a sterile iron spoon and stored in 50 ml sterile plastic tubes. The tubes were transported using dry ice to the lab, and stored at −80 °C for further analyses and treatment including DNA extraction. The genomic DNA was extracted from the samples using FastDNA SPIN (MP Biomedicals, Irvine, CA) according to the manufacturer’s instructions, and purified for library construction. The purified genomic DNA was subjected for metagenomic sequencing on an Illumina HiSeq2500 platform using paired-end 2 × 150 bp sequencing kit. The raw metagenomic reads were filtered to remove Illumina adapters, PhiX and other Illumina trace contaminants with BBTools Version 38.79, and low-quality bases and reads using Sickle (version 1.33; https://github.com/najoshi/sickle). The quality reads after filtering were assembled using metaSPAdes (version 3.10.1) with a kmer set of “21, 33, 55, 77, 99, 127”, and mapped to the corresponding assembled scaffolds using bowtie2^[55]^ (version 2.3.5.1) for sequencing coverage calculation. The coverage of a given scaffold was calculated using the MetaBAT2^[56]^ (version 2.12.1) script “jgi_summarize_bam_contig_depths”. For each sample, the scaffolds ≥ 2500 bp were binned using MetaBAT2 (version 2.12.1), with both tetranucleotide frequencies (TNF) and sequencing coverage of scaffolds considered. All the binned and unbinned scaffolds ≥ 1000 bp were uploaded to ggKbase (http://ggkbase.berkeley.edu/) for manual curation of genome bins based on GC content, sequencing coverage and taxonomic information of each scaffold^[57]^. The ggKbase genome bins were curated individually to fix local assembly errors using ra2.py as previously described^[58]^.

### Ribosomal Phylogeny

Genomes were searched for 16 ribosomal proteins^[59]^ using GOOSOS.py (https://github.com/jwestrob/GOOSOS) and the following HMMs: Ribosomal_L2 (K02886), Ribosomal_L3 (K02906), Ribosomal_L4 (K02926), Ribosomal_L5 (K02931), Ribosomal_L6 (K02933), Ribosomal_L14 (K02874), Ribosomal_L15 (K02876), Ribosomal_L16 (K02878), Ribosomal_L18 (K02881), Ribosomal_L22 (K02890), Ribosomal_L24 (K02895), Ribosomal_S3 (K02982), Ribosomal_S8 (K02994), Ribosomal_S10 (PF00338), Ribosomal_S17 (K02961), and Ribosomal_S19 (K02965). Ribosomal S10 model PF00338 was used in place of K02946 because the KOFAM model had a much lower hit rate than the other KOFAM models used.

Genomes containing at least 8 of these 16 proteins on a single scaffold were then used for further analysis. Retrieved protein sequences for each model were aligned individually using FAMSA^[90]^ and concatenated using the script Concatenate_And_Align.py (https://github.com/jwestrob/GOOSOS/blob/master/Concatenate_And_Align.py). The concatenated alignment was trimmed using trimal^[60]^ with the parameter -gt 0.1, keeping columns with fewer than 90% gaps. A guide tree was constructed using iQ-TREE^[61]^ with the LG+FO+R10 model, and the final phylogeny was constructed using iQ-TREE with the LG+C20+FO model^[63]^ and 1000 ultrafast bootstrap replicates^[62]^; the number of mixture model components was capped at 20 due to computational constraints.

### Rubisco

Rubisco large subunit sequences were identified using KOFAM model K01601 (*rbcL*) and PFAM model PF12338 (*rbcS*). Phosphoribulokinase sequences were identified using the PFAM model PF00485 (PRK). Rubisco subtype classification was performed using a phylogeny estimated using *Chloroflexi* supergroup *rbcL* hits as well as reference sequences from Jaffe et al. 2019^[64]^ as well as Banda et al. 2020^[13]^. Phylogeny estimation was performed using iQ-TREE, using the LG+FO+G4 model as well as the ultrafast bootstrap approximation.

Sequences classified as Form I, Form II (outgroup), Form I’, or Form I-⍺ were then extracted from this dataset. Sequences corresponding to form I-⍺ were then searched against ggKbase using BLASTP^[65]^, and sequences with greater than 95% identity from unbinned metagenomic contigs were then added to the sequence dataset. These protein sequences, as well as the previously classified Form I, Form II, Form I’, and Form I-⍺ sequences, were then used to build the phylogeny displayed in **Figure 3**. This phylogeny was estimated using iQ-TREE with the LG+FO+G4 model as well as the ultrafast bootstrap approximation. Genome context diagrams were then generated using Clinker^[66]^.

### Hydrogenases

Hydrogenase large subunit sequences were identified using the procedure outlined in Matheus-Carnevali et al. 2019^[67]^ using custom HMMs (attached, supplemental). *Chloroflexi* supergroup genomes were searched using the NiFe group 123, NiFe group 4, and FeFe HMMs, then classification was performed using a phylogeny built with these HMM hits and references from Matheus-Carnevali et al. 2019 and HydDB^[68]^. Hydrogenase phylogenies were constructed using iQ-TREE with the LG+FO+R models. Genome context diagrams in Figure **6** were generated using Clinker^[66]^.

Classification was then further refined by manual inspection of hydrogenase loci to ensure the presence of small subunit proteins as well as expected electron transfer and maturation machinery for each hydrogenase subtype.

The presence of genes encoding the catalytic subunit of hydrogenases was confirmed by phylogenetic analysis using references from Greening et al.^[8]^ (Fig. 6, Supp. Fig. S2, S3). Furthermore, visual inspection of hydrogenase gene clusters was performed and if at least the small subunit was not found in the vicinity of the large subunit, the genome was not included in hydrogenase counts. A combination of KOfam and Pfam annotations was used to determine the presence of any given gene cluster (Supplementary Table S3), although due to the less restrictive cutoffs for Pfam HMMs, the Pfam annotations were often used. Alternative annotations for the same genes were also taken into consideration in some cases (e.g., FeFe group C hydrogenases). The presence of a maturation protease in the vicinity of the hydrogenase genes, as well as multiple other hydrogenase expression/formation proteins, were considered as evidence of hydrogenase presence.

### Multiheme Cytochromes

Proteins from *Chloroflexi* supergroup genomes were searched for CxxCH heme-binding motifs using a regular expression in Python. Proteins with more than 20 such motifs were classified as multiheme cytochromes for the purpose of this analysis.

### Sulfur Cycle

———Sulfur cycling genes were identified using hits to KOFAM HMMs corresponding to kegg modules M00176 (Assimilatory sulfate reduction), M00596 (Dissimilatory sulfate reduction) and M00595 (Thiosulfate oxidation). Models used to search for genes involved in DMSP metabolism include K07306 (*dmsA*), K00184/K07307 (*dmsB*), K00185/K07308 (*dmsC*), K16964 (*ddhA*), K16965 (*ddhB*), K16966 (*ddhC*), K16967 (*dmoA*), and K17285 (SELENBP1).

### CAZys

Carbohydrate-Active enZymes (CAZys) were identified using dbCAN^[69]^ HMMs using an evalue threshold of 1e-15. CAZy counts were normalized by the size of the genome in mega-basepairs.

### CRISPR-Cas Systems

CRISPR-Cas loci were identified and counted using CRISPRcasIdentifier^[70]^ using default parameters. CRISPR repeats were identified using minced^[87]^.

### Identification of Genes Involved in the Nitrogen cycle

Genomes were classified as having functional nitrogenase if those genomes contained hits to nitrogenase alpha (K02586) and beta (K02591) subunits as well as the catalytic subunit nifH (K02588). *nxrAB* loci were identified using the *nxrAB* HMMs provided in Anantharaman et al. 2016^[6]^ using provided bitscore cutoffs, and scaffolds containing hits to both HMMs on the same scaffold were classified as *nxrAB*. N_2_O reduction via *norBC* was searched for using KOFAM models K04561 and K02305, with only genomes containing hits to both HMMs considered valid. Other multi-gene systems searched for via this method are *nirBD* (K00362 and K00363), *napAB* (K02576 and K03568), *nasAB* (K00372 and K00360), and *nrfAH* (K03385 and K15876), which similarly required both HMMs to have hits in the same genome to identify a functional system. Other nitrogen gene markers used are *nirK* (K00368), *nirS* (K15864), *nosZ* (K00376), *narB* (K00367), and *nirA* (K00366).

The *narGHIJ* complex was detected using the KOFAM models *narH/narY/nxrB* (K00371) and *narI/narV* (K00374); the catalytic subunit, *narG*, lacks a KOFAM HMM model and the TIGRFAM narG HMM (from TIGRFAM, Karthik’s Sulfur oxidation paper) was selected as an alternative model. Of those scaffolds containing the catalytic subunit *narG*, 73 had at least two of *narH, narI*, or *narJ*, and are considered complete for the purposes of this analysis.The presence of *narGHIJ* systems outside the clades in which it was detected by this analysis cannot be entirely discounted as systems with divergent *narG* sequences may exist that the TIGRFAM model does not capture with default cutoffs.

### Bacteriochlorophyll Biosynthesis Pathway Gene Identification

———Bacteriochlorophyll biosynthesis pathway genes were identified using KOFAMscan with provided bitscore cutoffs. Selected models include *bchI* (K03405), *bchE* (K04034), *bchD* (K03404), *bchH* (K03403), *bchZ* (K11335), *bchC* (K11337), *bchK* (K13605), *bchX* (K11333), *bchY* (K11334), *bchG* (K04040), *bchP* (K10960), *bchN* (K04038), *bchB* (K04039), *bchL* (K04037), *bciC* (K21058), *bchF* (K11336), *bchJ* (K04036), *acsF* (K04035), *bchM* (K03428), NYC1 (K13606), and *fmoA*/bacteriochlorophyll A protein (K08944). Where PFAM HMM models specific to a particular bacteriochlorophyll synthesis gene were available, the union of the corresponding PFAM and KOFAM HMMs were taken to represent hits to that particular gene, including *bchJ* (PF02830/V4R), *bchL* (PF02043/Bac_chlorC), *bchF* (PF07284/BCHF), *fmoA*/bacteriochlorphyll A protein (PF02327/BChl_A), and *bchM* (PF07109/Mg-por_mtran_C).

Chlorosome genes were defined as hits of greater than or equal to 40% identity to representative chlorosome genes from *Chloroflexus aurantiacus* available in uniprot (*csmA, csmM, csmN, csmO*).

Sequences for photosystem reaction center II subunits *pufL* and *pufM* within the *Chloroflexi* supergroup were obtained by searching with pfam model PF00124 (*Photo_RC*) and applying the model-designated GA cutoff; genomes with two hits, one for each subunit, were considered to have the *pufLM* complex. Proteins containing two domain-level hits to PF00124 and approximately twice the length of *pufL* were considered *pufLM* fusion events.

### Reductive Dehalogenases

Reductive dehalogenase enzymes were searched for using the PFAM model PF13486 (Dehalogenase) as well as KOFAM models K01560, K01563, and K01561. The count of reductive dehalogenase enzymes per genome (**Fig. 2**) is defined as the union of all such hits per genome.

## Data Availability

All supplementary data, including nucleotide and protein fasta files for each genome in the dataset and associated annotations, for this project is available at https://figshare.com/projects/Chloroflexi_Supergroup/120267.

## Supplementary Figures

**Supplementary figure S1.**
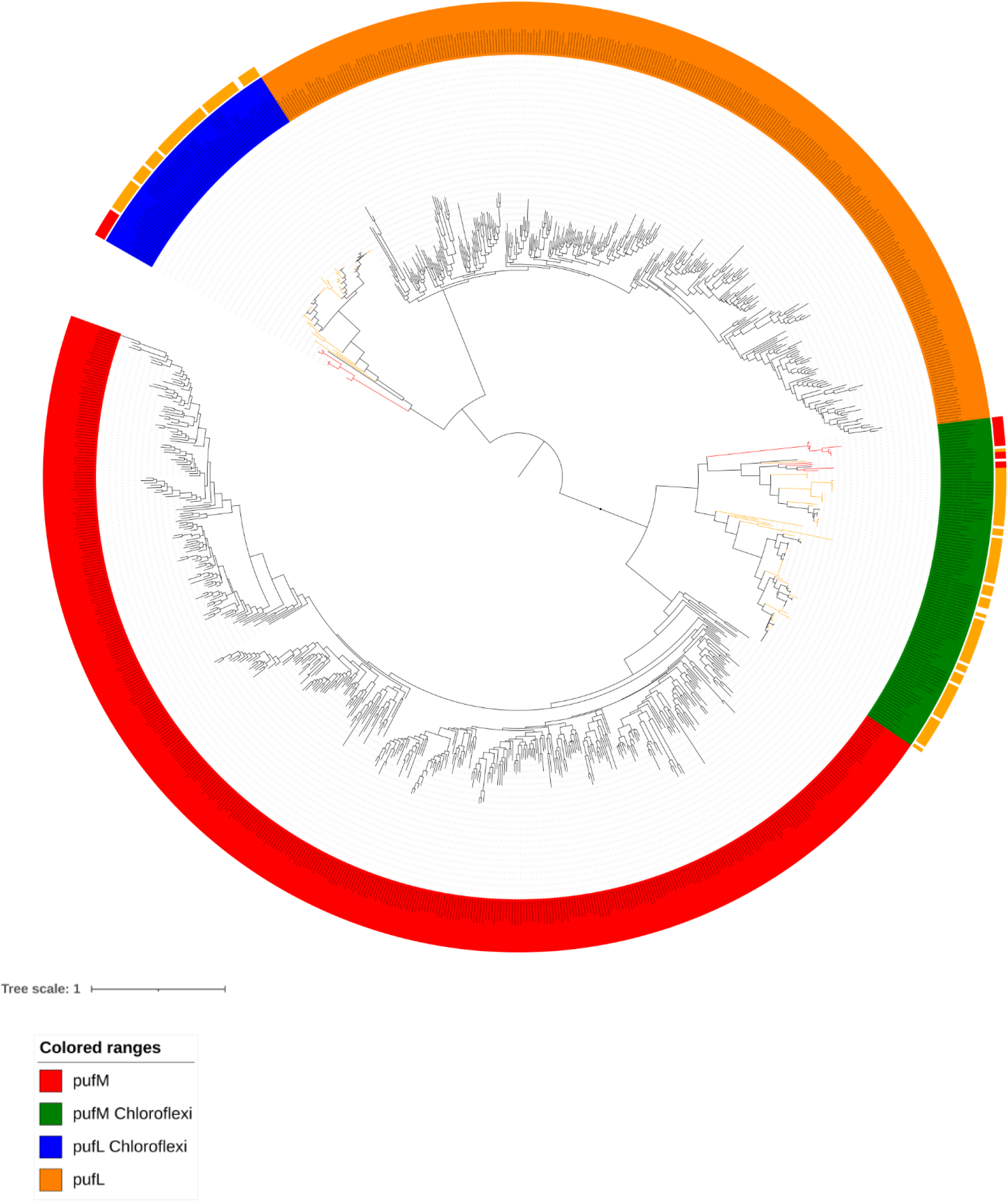
Phylogeny of type II photosystem reaction centers *pufLM* with reference sequences from Uniprot. Sequences with decorations on the outer ring indicate *pufLM* sequences from the *Chloroflexi* supergroup genome dataset; red decorations indicate *Anaerolinea* and orange indicate *Chloroflexia*.

**Supplementary figure S2.**
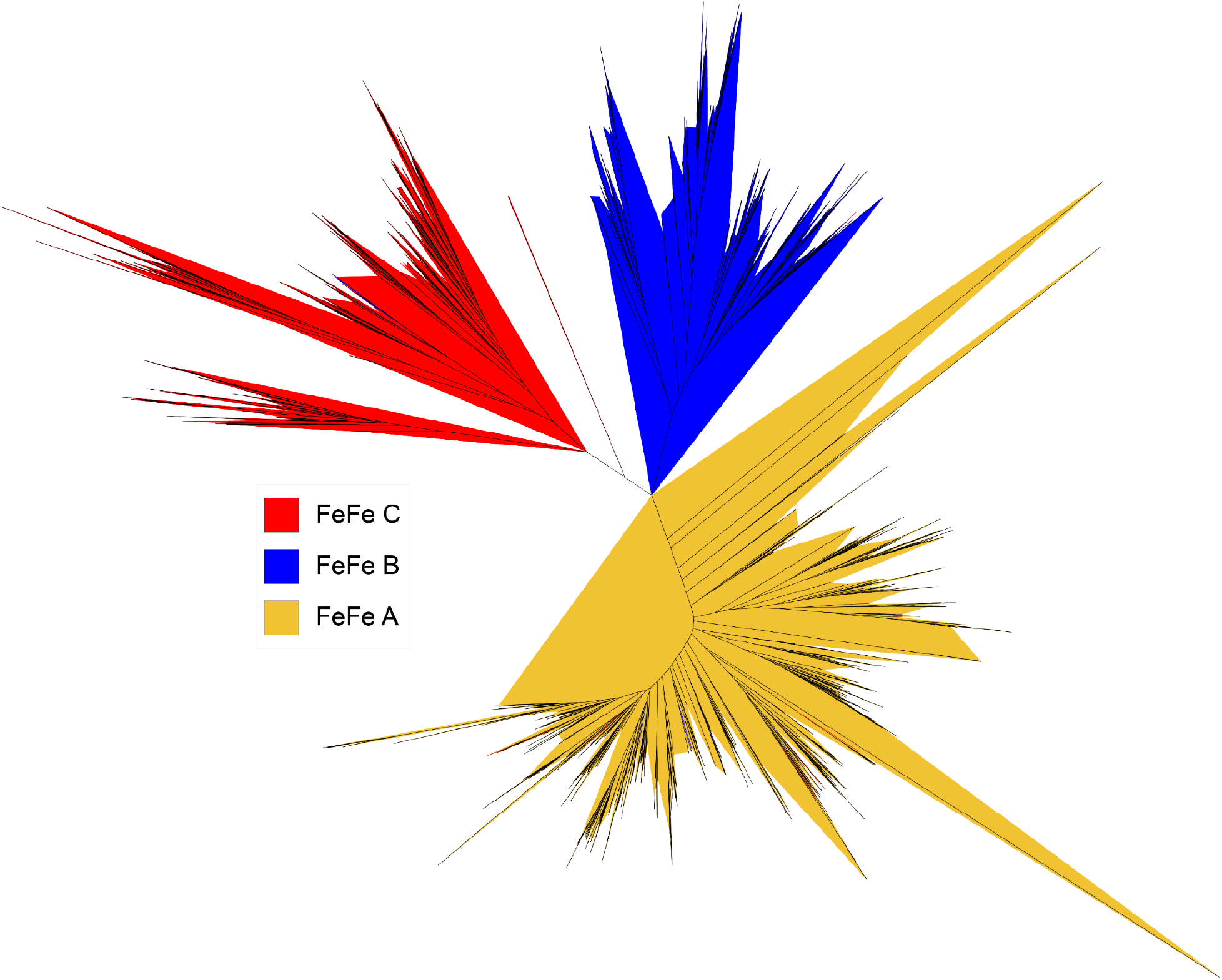
Phylogeny of FeFe hydrogenases in the *Chloroflexi* supergroup including references from HydDB and Matheus Carnevali et al. 2019.

**Supplementary figure S3.**
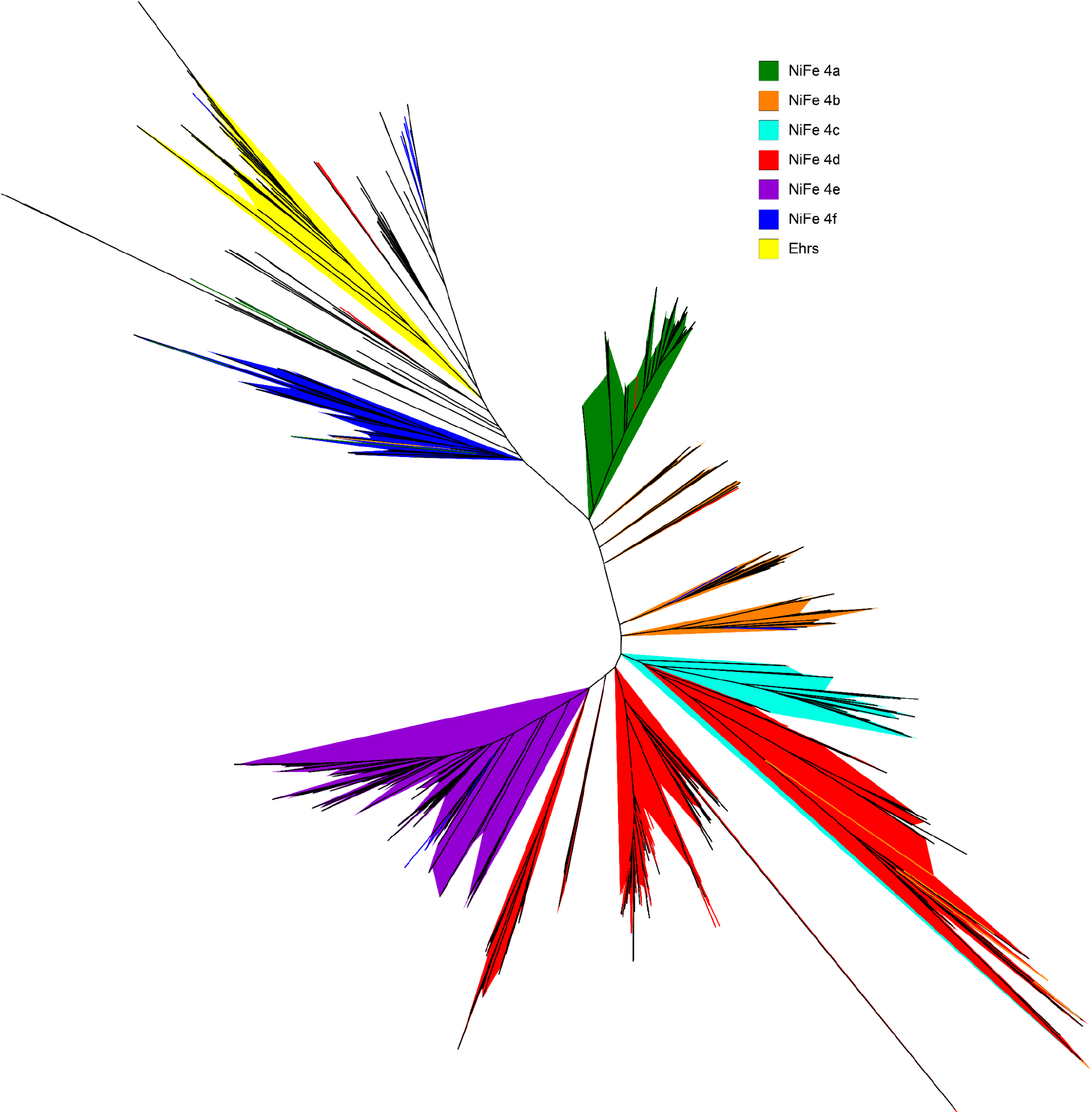
Phylogeny of NiFe group 4 hydrogenases in the *Chloroflexi* supergroup including references from HydDB and Matheus Carnevali et al. 2019.

**Supplementary figure S4.**
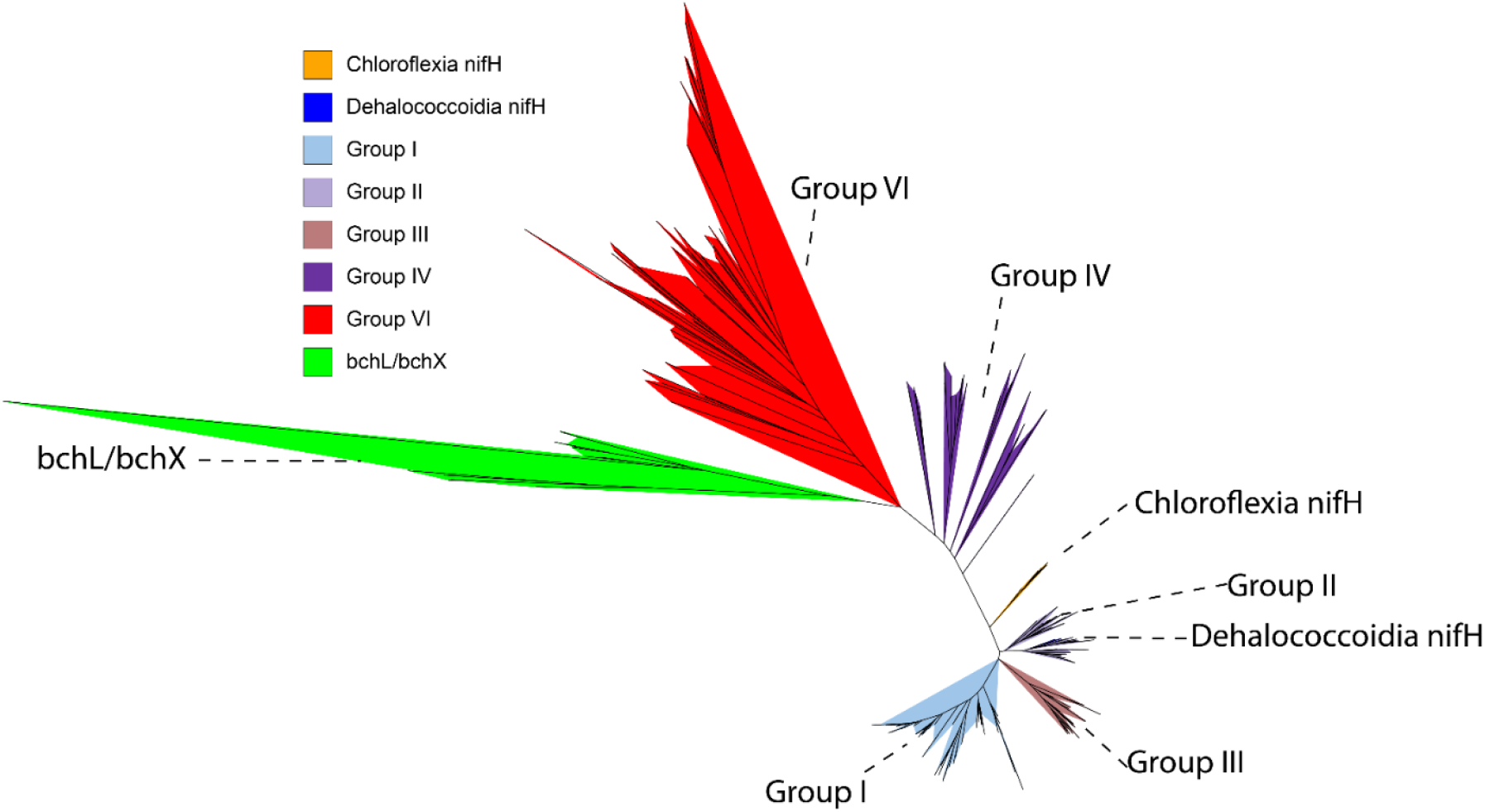
Phylogeny of *nifH* sequences with references from Uniprot and Meheust et al. 2020.

**Supplementary figure S5.**
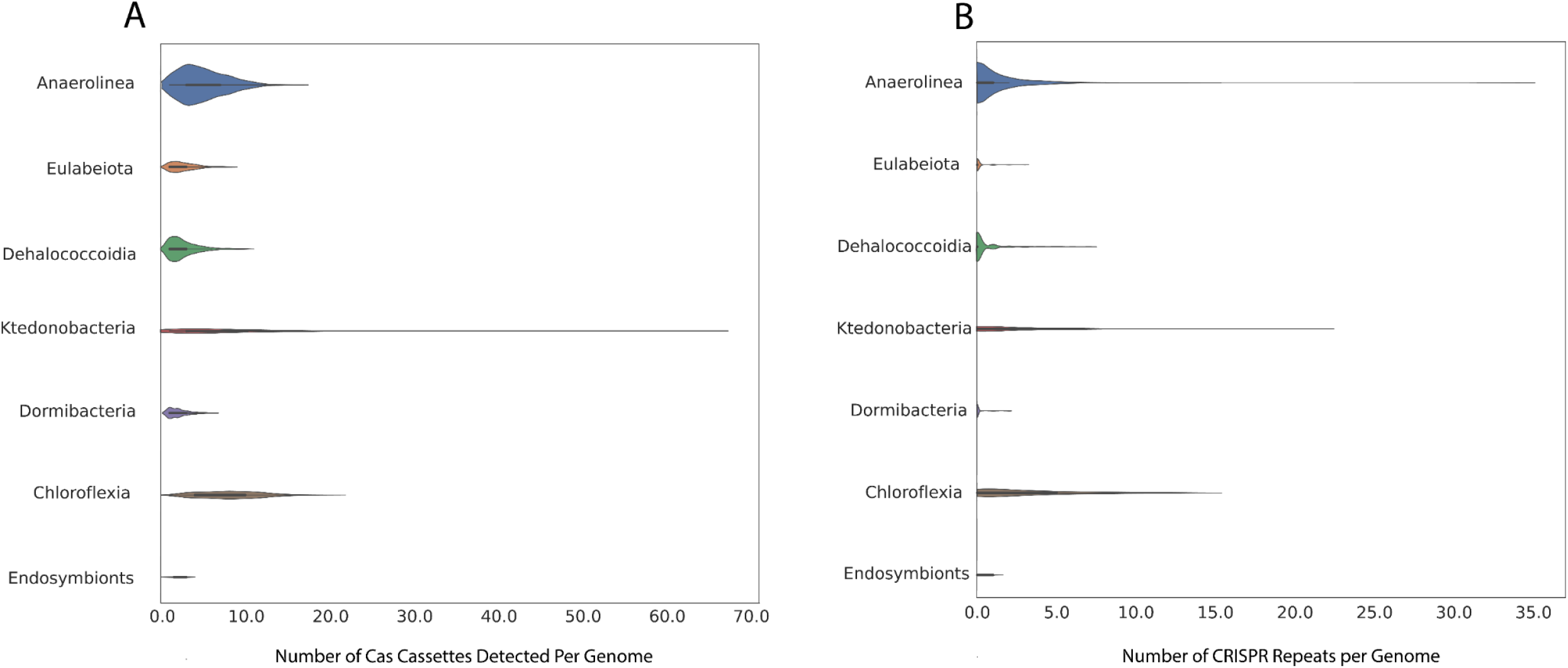
Violin plot of number of CRISPR loci by phyletic group within the *Chloroflexi* supergroup.

